# 2-Arachodonoylglycerol-mediated endocannabinoid signaling modulates mechanical hypersensitivity associated with alcohol withdrawal in mice

**DOI:** 10.1101/2022.02.15.480609

**Authors:** Amanda Morgan, Danielle Adank, Keenan Johnson, Emily Blunt, Sachin Patel

## Abstract

Alcohol use disorder (AUDs) commonly co-occurs in patients with chronic pain, and a major barrier to achieving abstinence and preventing relapse is the emergence of hyperalgesia during alcohol withdrawal. Elucidating novel therapeutic approaches to target hyperalgesia associated with alcohol withdrawal could have important implications for the treatment of AUD. Here we examined the role of 2-arachidonoylglycerol (2-AG)-mediated endocannabinoid (eCB) signaling in the regulation of hyperalgesia associated with alcohol withdrawal in mice and tested the hypothesis that pharmacological augmentation of 2-AG signaling could reduce hyperalgesia during withdrawal. After 72 hours of withdrawal from a continuous access two-bottle choice drinking paradigm, male and female mice exhibited increased mechanical but not thermal hypersensitivity, which normalized by 7 days. This effect was reversed by pretreatment with the monoacylglycerol lipase (MAGL) inhibitor JZL184, which elevates levels of 2-AG. The effects of JZL184 were prevented by coadministration of either a CB1 or CB2 antagonist. Inhibition of the 2-AG synthetic enzyme diacylglycerol lipase (DAGL) with DO34 exacerbated mechanical hypersensitivity during alcohol withdrawal, causing an earlier onset and persistent hypersensitivity even one week into withdrawal. Our findings demonstrate the critical role of 2-AG signaling in the bidirectional regulation of mechanical sensitivity during alcohol withdrawal, with enhancement of 2-AG levels reducing sensitivity, and inhibition of 2-AG synthesis exacerbating sensitivity. These data suggest 2-AG augmentation could represent a novel approach to the treatment of alcohol withdrawal-associated hyperalgesia and AUD in patients with comorbid pain disorders.

## INTRODUCTION

The lifetime prevalence of alcohol use disorders (AUDs) is ~30% in the U.S. population [1], with an estimated 15.5 million U.S. adults suffering from AUDs [2]. Disorders of acute and chronic pain often co-occur with AUDs, with chronic pain affecting an estimated 114 million adult Americans [3, 4]. Alcohol has analgesic properties, with 38% of heavy drinkers reportedly drinking to treat pain [5]. Drinking to cope with physical pain can be effective initially due to the analgesic properties of alcohol [6]. However, the analgesic effects of alcohol are brief and dose-dependent, with tolerance emerging relatively quickly [6, 7]. These pharmacological properties of alcohol can lead to physiological dependence on alcohol for pain relief, contributing to negative reinforcement driven alcohol use [3, 8]. Importantly, neurotoxic effects of alcohol can result in alcoholic neuropathy after chronic use. Furthermore, alcohol withdrawal is associated with the emergence of hyperalgesia [9–13] and allodynia which can persist after weeks or months of abstinence [10]. Moreover, this effect is seen in patients who are alcohol dependent but report no preexisting chronic or acute pain conditions. For example, pain-free patients may develop a new heightened sensitivity to painful stimuli when they undergo withdrawal from alcohol after prolonged use [3, 10]. This effect has been well documented in preclinical studies where alcohol withdrawal-induced hyperalgesia is extensively reported in rodents [14–32].

Major barriers to achieving abstinence and preventing relapse include the emergence of negative affect and increased pain sensitivity during alcohol withdrawal [3, 33]. Effective pharmacological treatments targeting the interdependence between alcohol and pain are currently limited. For example, Gabapentin is used to treat both acute [34–39] and chronic pain conditions [40–45], and has recently emerged as a possible pharmacotherapeutic for AUDs [46–52]. The Federal Drug Administration has approved the drugs disulfiram, acamprosate, and naltrexone to treat AUDs; however, clinical data indicate these medications are only partially effective and have high rates of relapse upon discontinuation of the drugs [53]. None of the medications approved to treat AUDs are useful for treatment of pain associated with alcohol withdrawal. These data underscore the high need to identify new potential therapeutic approaches for the treatment of alcohol withdrawal associated hyperalgesia, which could facilitate abstinence by reducing negative reinforcement driven alcohol use in treatment-seeking individuals.

One emerging target for the treatment of pain is the endogenous cannabinoid (eCB) system, which acts throughout neural nociceptive system to produce antinociceptive and analgesic effects [54, 55]. The eCBs 2-Arachidonoylglycerol (2-AG) and N-arachidonoylethanolamine (AEA) exert antinociceptive effects in rodents through activation of CB1 and CB2 receptors [56]. These eCBs are also implicated in setting nociceptive thresholds and regulating tonic inhibition of pain responses [56, 57]. Pharmacological inhibition of the major 2-AG catabolic enzyme monoacylglycerol lipase (MAGL) results in increased brain and peripheral levels of 2-AG and exerts analgesic and anti-allodynic effects of in a variety of preclinical models via CB1 and CB2 receptor [58–66].

Here we utilized a continuous access two bottle choice alcohol drinking paradigm combined with pharmacological approaches to test the hypothesis that 2-AG signaling regulates pain sensitivity during alcohol withdrawal. We report that pharmacological augmentation of 2-AG during alcohol withdrawal reverses mechanical hypersensitivity associated with alcohol withdrawal in mice via actions at both CB1 and CB2 receptors. Conversely, pharmacological 2-AG depletion worsens and extends the time course of hyperalgesia observed during alcohol withdrawal, suggesting a critical role for endogenous 2-AG to counteract hyperalgesia during withdrawal. These data provide new insight into the mechanisms underlying alcohol withdrawal associated hyperalgesia and suggest 2-AG augmentation could represent a novel treatment approach of AUD in patients with comorbid pain disorders.

## METHODS AND MATERIALS

### Animals

C57BL/6J (Jackson Laboratory, ME) female and male mice were between 6-7 weeks old at the beginning of experiments. The animal care facilities at Vanderbilt University (Nashville, TN), housed mice in climate-controlled colony rooms, maintained at 21 ± 2°C, 30% ± 10% relative humidity on a 12L: 12D cycle, with lights on at 0600h. Food and water were provided *ad libitum*(LabDiet 5001; LabDiet) for the duration of the experiments. All behavior experiments were conducted during the light phase. All studies were carried out in accordance with the National Institute of Health Guide for the Care and Use of Laboratory Animals and approved by the Vanderbilt University Institutional Animal Care and Use Committee (#M1600213-01).

### Drugs

The MAGL inhibitor JZL184 (10 mg/kg^-1^, Cayman Chemical, MI, USA), CB_1_R inverse agonist Rimonabant (3 mg kg^-1^ APIChem, Hangzhou, Zhejiang, China), and CB_2_R inverse agonist AM630 (3 mg kg^-1^, Cayman Chemical, MI, USA) were prepared in DMSO and injected i.p.at 1 μl g^-1^ bodyweight. DAGL inhibitor DO34 (50 mg kg^-1^, Glixx Laboratories Inc., MA) was administered in a vehicle mixture of ethanol (Pharmco, KY, USA): kolliphor (Sigma-Aldrich, WI,): saline (Hospira, IL, USA) [1:1:18] was injected at a volume of 10 μl g^-1^ bodyweight. All drugs were administered 2 hours before behavior testing. 190 proof ACS/USP grade grain-derived EtOH (Pharmco, KY, USA) was used to for EtOH drinking solutions.

### Two-bottle choice EtOH drinking and withdrawal model

Mice were first acclimated to single-housed two-bottle choice (2BC) cages for 7 days, with two sippers and access to tap water only. For EtOH mice, one bottle of tap water was replaced with 3% EtOH in tap water for 4 days. On the 5^th^ day, the EtOH concentration increased to 7% for 7 days. On the 8^th^ day, the EtOH bottle concentration increased to 10% or 20% and were continues at this concentration until testing or forced abstinence. EtOH bottle placement (left or right sipper) was rotated weekly. For withdrawal tests, after 21 days of drinking in 2BC cages with either one bottle of 10% or 20% EtOH, the EtOH bottle was replaced with tap water. Control mice were housed in the same conditions for the duration of the experiment with two bottles of tap water. Mice undergoing EtOH withdrawal were kept physically separated from control mice in the facility to prevent social transfer of pain sensitivity, by use of adjoining enclosed housing rooms [19].

### Pain Assays

Mechanical punctate sensitivity was assessed by manual application of Von Frey filaments of varying forces (0.4–4.0g) using the “ascending stimulus” method [67]. Mice were tested in groups of four in a plexiglass testing rack, which had 4 clear plexiglass experimental cubbies boxes. Because mice were single housed for a protracted period, the interior walls of each cubby were covered with a thin piece of dark plastic, so the mice could not see each other during habituation and testing. Each mouse had left hind paw tested first, with increasing grams force monofilaments applied until a positive response was observed. ‘Positive response’ included of paw withdrawal, paw licking, or shaking, either during application or immediately after the filament was removed. The grams force of the von Frey filament that elicited a positive response was designated as the mechanical withdrawal threshold for that trial. The trial was repeated once again for the left hind paw, with monofilaments applied in ascending grams force until a positive response. Next, the second mouse on the apparatus was tested for two trials on left hind paw, followed by third mouse in the apparatus and the fourth. Then, the same two-trial procedure was repeated for each mouse using the plantar surface of each right paw. Averaged paw withdrawal thresholds are reported [68] Thermal sensitivity was assessed with the hot plate test, performed as previously described [69], with mice placed onto a flat surface pre-heated to 50°C within an open Plexiglas tube. Latency to the first positive response was recorded, with a cutoff time of 40 seconds. Positive responses included hind paw shaking or hind paw licking, or mouse jumping. Latency in seconds to positive response or reaching cutoff time with no positive responses for one trial per mouse is reported.

### Irritability-like behavior tests

To measure “irritability”, we used the Bottle Brush Test for mice [70–72]. After 30 minutes of acclimation to the testing room, mice were placed alone in their open home cage. Two cameras were aimed at the clear cage, one from a lateral side view and one overhead. The experimenter used a bottle brush – a small white plastic brushed with a rounded top of protruding plastic bristles, used to clean small bottles. Briefly, mice were lightly “attacked” by the brush (touched by the bristles of the brush and followed around by the brush) in the following sequence:

1. Brush (in rotation) approaching the mouse from the end (starting position) of the cage.
2. Brush (in rotation) touching the whiskers of the mouse.
3. Brush (in rotation) returning to the starting position in the opposite end of the
4. Brush (in rotation) at the starting position.
5. Brush (no rotation) at the starting position.

Each mouse was attacked by the bottle brush 20 times per session. Videos were coded using ANVIL video coding software [73] for the following positive responses: (1) mouse was climbing cage wall during the bottle brush attack, (2) digging, (3) running away/escaping the bottle brush, (4) freezing in response to the bottle brush attack, or (5) exploring the bottle brush as it was touching them. Using the ANVIL time coding function, the during of each positive response was recorded. The total time for each positive response during 20 bottle brush attacks is reported for each mouse.

### Anxiety-like behavior tests

Elevated Plus Maze (EPM), Open Field Test (OFT) and Light-Dark Box (LD) were performed exactly as previously described [74].

The EPM had two wall-free open-arms (30 × 10 cm; light illuminance ~100-115 lux), and 2 walled (“closed”) arms (30 × 10 × 15 cm; ~20-25 lux) anchored to a square 5 × 5 cm open center. To begin the test and trigger video tracking, mice were placed in the center, facing an open arm. Maze was made of white acrylonitrile butadiene styrene plastic and elevated 47 cm off the ground. Each mouse explored the EPM for 5 minutes per test, with ANY-maze (Stoelting, Wood Dale, Illinois) video tracking software used to monitor and analyze behavior during testing.

The OFT used a square sound-attenuating chamber with clear plexiglass walls (27.9 × 27.9 × 20.3 cm; MED-OFA-510; MED Associates, St. Albans, Vermont) contained within a white sound-attenuating chamber box. Mice were placed in the center of the open field chamber to begin the test and recorded for 5 minutes, with beam breaks from 16 infrared light beams measuring movement and position with Activity Monitor v5.10 (MED Associates) software. The center of the OFT was designated as a square of the innermost 50% of the OFT arena floor, with the remainder considered the perimeter. The chamber was illuminated at ~200 lux and white noise was present at ~60 dB.

The LD box test used a black insert (Med Associates ENV-511; made of IRT to allow infrared beam transmission) was placed into a chamber with clear plexiglass walls (27.9 × 27.9 × 20.3 cm; MED-OFA-510) to split the chamber into equal halves light (~350-400 lux) and dark (<5 lux) divisions. Beam breaks from 16 infrared beams were recorded Activity Monitor v5.10 (MED Associates) to monitor position and behavior during the 10-min test. White noise was present at ~60 dB.

## RESULTS

### Hyperalgesia during EtOH withdrawal

We first investigated the effects of withdrawal from 10% EtOH on nocifensive behaviors (Fig.1). Mechanical and thermal sensitivity were measured 24 hours, 72 hours, and one week into withdrawal (Fig.1A). Compared to water drinking controls, EtOH withdrawal mice showed increased mechanical hypersensitivity, with significantly lower paw withdrawal thresholds on the Von Frey filament test at 72 hours into withdrawal. However, no significant effect of EtOH withdrawal was seen at the earlier withdrawal timepoint of 24 hours, or at the late timepoint of one-week post EtOH withdrawal (Fig.1B). In contrast, there was no difference between alcohol withdrawal and water-drinking controls on the hot plate test of thermal sensitivity at any time point (Fig.1C). A separate cohort of mice were tested on the Von Frey filament and hot plate tests after three weeks stable drinking on 10% EtOH in 2BC, i.e. not in withdrawal (Fig. 1D). Mice drinking 10% EtOH in 2BC were not different from water controls on either Von Frey (Fig. 1E) or hot plate (Fig. 1F) thresholds. These data indicate that alcohol withdrawal is associated with a timed-dependent emergence of mechanical, but not thermal, hypersensitivity, and that this effect is not observed during alcohol drinking *per se*, and thus selective to the withdrawal state.

**Figure 1.**
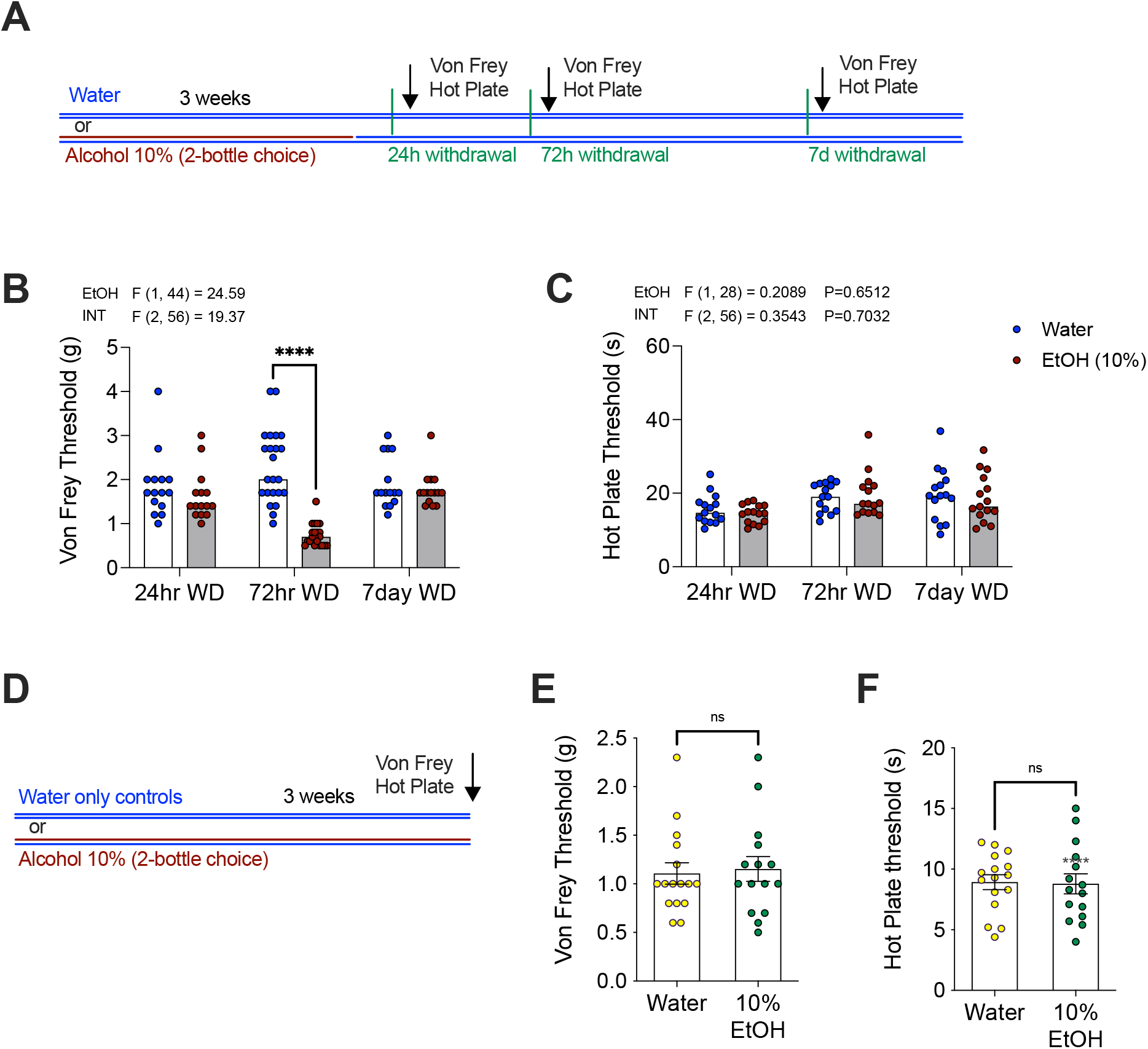
Alcohol withdrawal produces transient mechanical but not thermal hypersensitivity. **(A)** Schematic of 21-day 2BC EtOH drinking paradigm and behavioral testing at withdrawal time points. (**B)** Mechanical hypersensitivity is seen only at the 72-hour EtOH withdrawal time point, with reduced Von Frey threshold compared to water-drinking controls. (**C)** No effects on thermal sensitivity were observed at any withdrawal timepoint measured by the Hot Plate test. (**D)** Schematic of pain testing during drinking 10% EtOH or water-only. No effect on (**E**) Von Frey or (**F**) Hot Plate threshold was observed during drinking. F-scores on graphs represent main effects or interactions from Two-Way ANOVAs. **** p<0.0001 via post-hoc Holm-Sidak pairwise comparisons. Data presented as individual female mice and Mean ± SEM. NS; not significant.

### JZL184 reverses mechanical hypersensitivity associated with EtOH withdrawal

We next conducted several independent experiments to determine if pharmacologically increasing 2-AG levels affected the increased mechanical hypersensitivity observed during EtOH withdrawal. First, baseline Von Frey threshold was determined (Fig. 2A) prior to initiation of the 2BC model. After ramping up from 3%-10% EtOH, mice drank 10% EtOH for three weeks, and were tested using the Von Frey assay at 72 hours into withdrawal (Fig. 2A). EtOH withdrawn mice treated with vehicle had significantly reduced Von Frey thresholds compared to their own baselines whereas no changes in Von Frey threshold were detected in water control mice across these time points. (Fig. 2C). Beginning immediately after 72hr withdrawal Von Frey testing was completed, the EtOH mice were placed back on 2BC 10% EtOH for two additional weeks. After two weeks of additional drinking, EtOH was again removed for mechanical sensitivity testing at 72 hours withdrawal. This time, water mice and EtOH mice were pretreated with MAGL inhibitor JZL184 (10mg kg^-1^) to determine if increasing 2-AG levels could reverse hyperalgesia at 72 hours of EtOH withdrawal, and if JZL184 has analgesic properties in control mice drinking water only (Fig. 2A). JZL184 normalized mechanical hypersensitivity by significantly increasing Von Frey threshold in mice in EtOH withdrawal (Fig. 2B-C). The water-only group tested alongside the withdrawal mice showed no differences when tested at baseline, with vehicle, or with JZL184 pretreatment.

**Figure 2.**
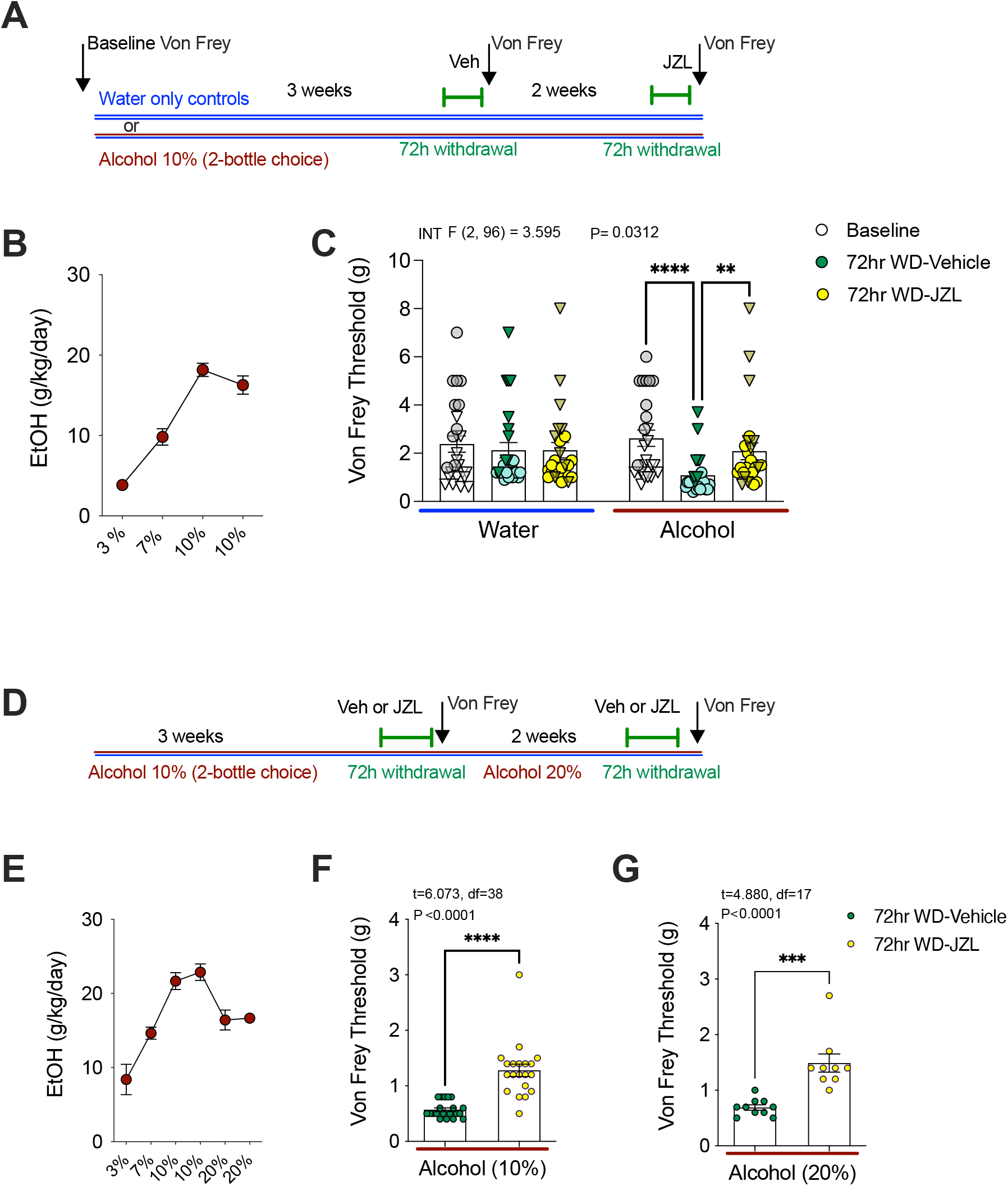
JZL184 reverses mechanical hypersensitivity at 72 hours EtOH withdrawal. **(A)** Schematic of experimental timeline. **(B)** Weekly EtOH intake of alcohol-drinking mice. **(C)** Von Frey threshold is significantly reduced at 72 hours EtOH withdrawal, which is prevented by JZL184. **(D)** Schematic experimental timeline. **(E)** Weekly EtOH intake of alcohol-drinking mice. **(F-G)** JZL184 reverses mechanical hypersensitivity at 72 hours EtOH withdrawal from 10% and 20% EtOH. F and t statistics shown in figures from 2-Away ANOVA interaction and unpaired t-tests. ** p<0.01, *** p<0.001, **** p<0.0001 via post-hoc Holm-Sidak pairwise comparisons or t-test. Data presented as individual mice (triangles males and circles females) and Mean ± SEM.

To confirm these findings using a between-subjects design, a separate cohort of 2BC EtOH drinking mice were treated with vehicle or JZL184 (10mg kg^-1^) and tested at 72 hours withdrawal from 10% EtOH (Fig. 2D). Consistent with previous experiment, JZL184 was able to significantly increase Von Frey thresholds relative to vehicle treatment at 72h into EtOH withdrawal (Fig. 2E-F). Immediately after Von Frey testing with vehicle or JZL184 pretreatment during 72h of withdrawal from 10% EtOH, mice were put back into 2BC cages and given one bottle of water and one bottle of 20% EtOH. After two weeks of drinking, access to 20% EtOH was removed, and Von Frey testing was gain conducted at 72h into withdrawal (Fig. 2D). Again, 72 hours after access to 20% EtOH was removed, mechanical hypersensitivity was seen in vehicle treated mice (Fig. 2G). There were no differences in the hyperalgesia triggered by removal of access from 20% EtOH (Fig. 2G) when compared to mechanical hypersensitivity seen 72 hours after removal of access from 10% EtOH (Fig. 2F, Fig. 2C, and Fig. 1C). Again, confirming out previous experiment, JZL184 was also able to reverse mechanical hypersensitivity 72h into withdrawal from 20% EtOH (Fig. 2G). These data provide compelling evidence that MAGL inhibition can reduce mechanical hypersensitivity induced by alcohol withdrawal.

### Hyperalgesia during EtOH withdrawal does not depend on concurrent increases in anxiety-like behavior

We considered the possibility that EtOH withdrawal-induced hyperalgesia could be considered one effect among several seen in the broader context of a general aversive behavioral response to alcohol withdrawal. To determine if mechanical hypersensitivity was an independent effect of EtOH withdrawal or was contingent upon an increase in anxiety-like behavior, the open field test was done during 72-hour withdrawal with a new cohort of 10% EtOH drinking mice. The EtOH withdrawal mice were no different in the total or center distance travelled in the open field test but spent more time in the center of the apparatus than water-drinking controls (Fig. 3). After this, the light-dark box and elevated-plus maze tests were conducted after mice were put back on EtOH for two weeks to achieve stable drinking levels, then EtOH was removed for 72 hours (Fig. 3A). Mice in EtOH withdrawal did not exhibit increase anxiety-like behavior on any measure. The lack of differences between groups indicates the mechanical hypersensitivity see in EtOH withdrawal groups is not associated with increases in anxiety-like behaviors at the 72 hour timepoint.

**Figure 3.**
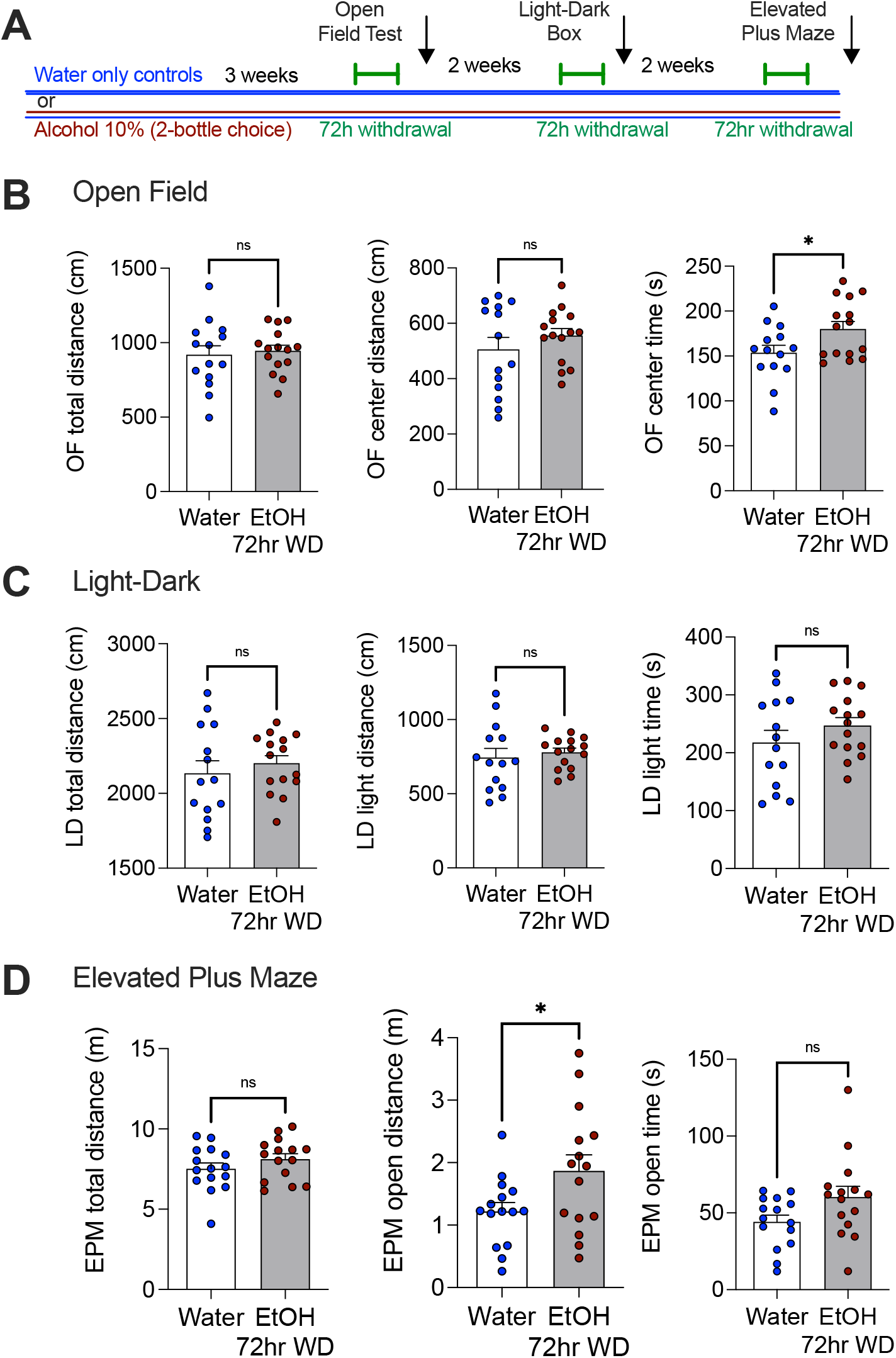
EtOH withdrawal does not increase anxiety-like behaviors. Schematic of experimental timeline. **(B)** Effects of 72-hour EtOH withdrawal on Open Field test. **(C)** Effects of 72-hour EtOH withdrawal on Light-Dark test. **(D)** Effects of 72-hour EtOH withdrawal on Elevated Plus Maze. ** p<0.01, via t-test. Data presented as individual female mice and Mean ± SEM.

### EtOH withdrawal induces irritability behavior, which is not prevented with MAGL inhibition

In the next experiment, mice in EtOH withdrawal were tested to determine if they expressed concurrent states of irritability-like behavior during EtOH withdrawal. Compared to water-only controls, EtOH withdrawal mice engaged in fleeing, but not digging, behavior significantly more often in the bottle brush irritability test (Fig. 4A). Furthermore, treatment with JZL-184 (10mg kg^-1^) was not able to prevent increases in irritability behavior in a separate cohort of EtOH drinking mice in 72-hour withdrawal (Fig. 4B). EtOH withdrawal thus results in irritability and mechanical hypersensitivity. This lack of sensitivity to JZL184 suggests the neural mechanisms underlying mechanical hypersensitivity triggered by EtOH withdrawal and irritability-like behaviors are likely dissociable.

**Figure 4.**
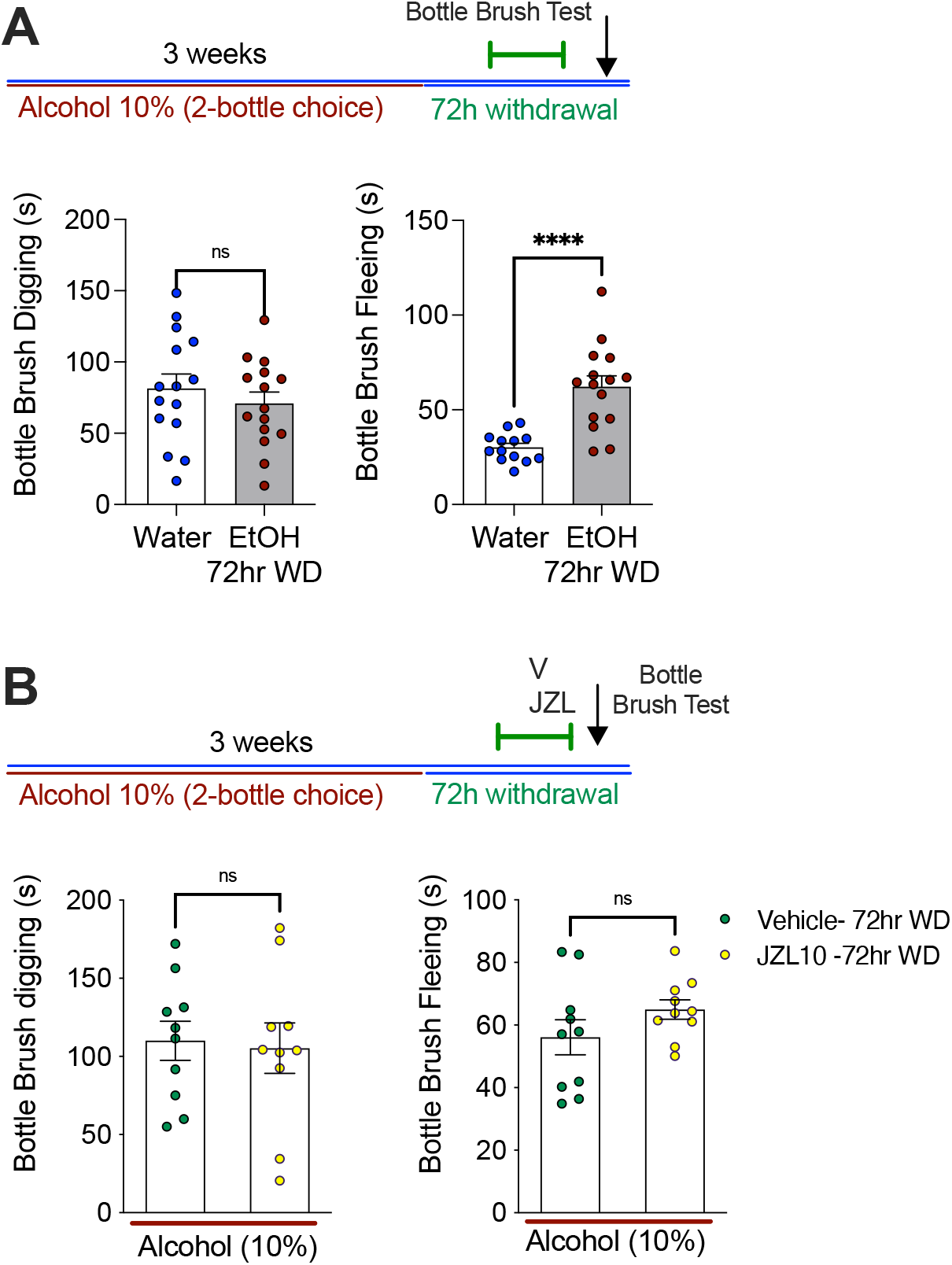
EtOH withdrawal increases irritability behavior, which is not prevented by JZL-184. A. Schematic of 2BC drinking with irritability behavior tested at 72 hours EtOH withdrawal. (Left) 72-hour EtOH withdrawal had no effect on digging behavior in the bottle brush test, but (right) significantly increased fleeing behavior. B. A separate cohort was tested for irritability at 72-hours withdrawal and given vehicle or JZL-184. (Left) JZL-184 had no effect on digging or (right) fleeing in the bottle brush test. **** p<0.0001 via t-test. Data presented as individual female mice and Mean ± SEM.

### Both CB1 and CB2 receptor antagonists prevent the anti-hyperalgesic effects of MAGL inhibition

Because JZL184 reverses mechanical hypersensitivity during EtOH withdrawal, we next sought to further characterize the pharmacological mechanisms underlying these effects. To determine if the anti-hyperalgesic properties of 2-AG are mediated through CB_1_ or CB_2_ receptors, mice underwent 72 hours of EtOH withdrawal and were treated with JZL184 alone, or JZL184 in combination with CB_1_R and CB_2_R antagonists (Fig. 5). Mechanical hypersensitivity returned when JZL184 was co-administered with either the CB_1_R antagonist Rimonabant (3mg kg^-1^), or the CB_2_R antagonist AM-630 (3mg kg^-1^). The antinociceptive effect of JZL184 was completely blocked by both CB receptor antagonists (Fig. 5A). No differences in Von Frey threshold were seen in control mice that only drank water and were treated with Rimonabant (3mg kg^-1^), or AM-630 (3mg kg^-1^), two hours before the Von Frey testing (Fig. 5B). This implies the involvement of CB_1_ and CB_2_ receptors in the anti-hyperalgesic effects of JZL184.

**Figure 5.**
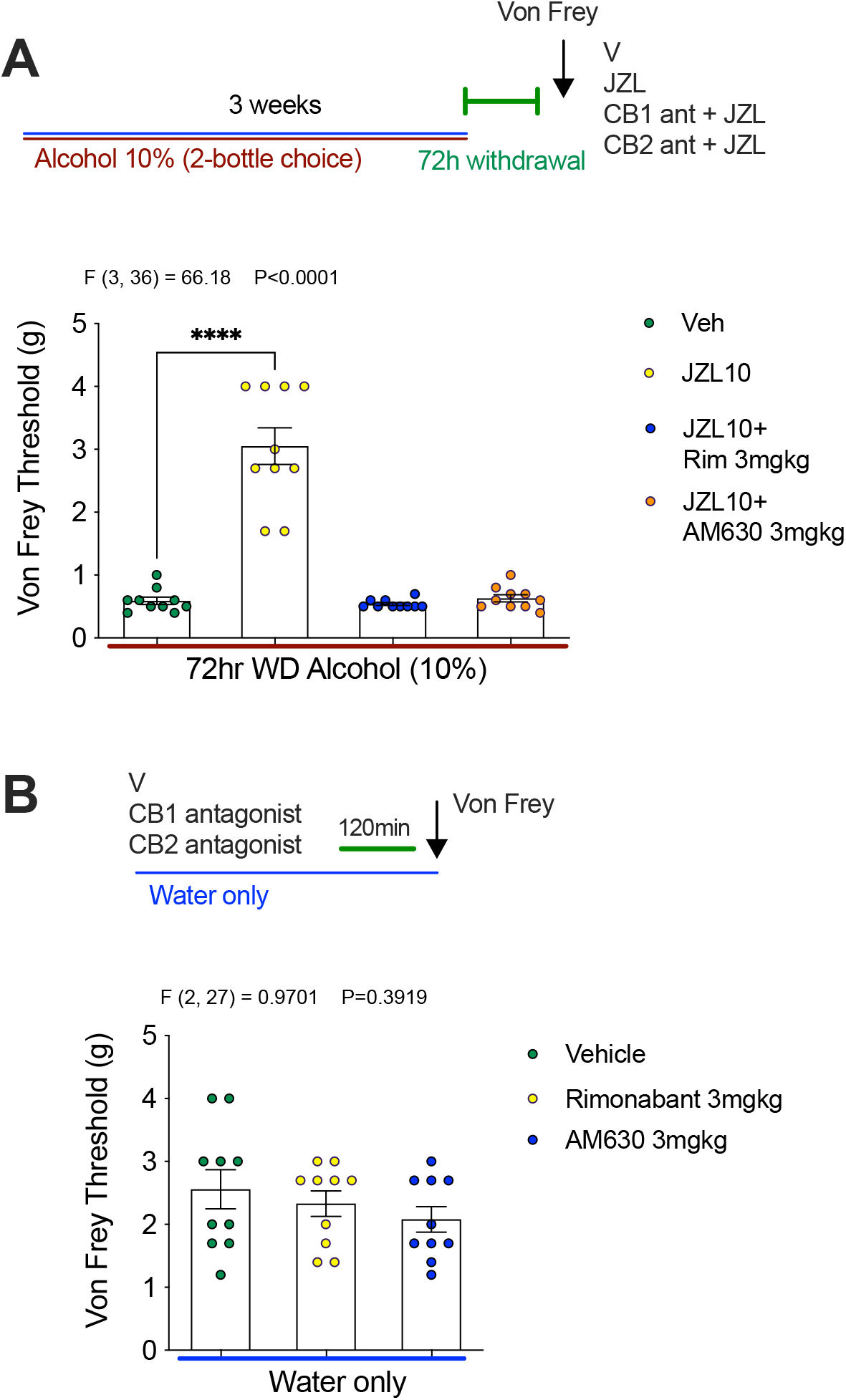
CB1 and CB2 receptor antagonists prevent the anti-hyperalgesic effects of JZL184. **(A)** Both Rimonabant and AM630 prevent the anti-hyperalgesic effects of JZL184. **(B)** Neither Rimonabant nor AM630 affect mechanical sensitivity thresholds in water drinking mice. F-scores on graphs represent main effects of ANOVA. ****p<0.0001 via post-hoc Holm-Sidak pairwise comparisons. Data presented as individual female mice and Mean ± SEM.

### The DAGL inhibitor DO34 enhanced mechanical hypersensitivity during EtOH withdrawal

The lowered mechanical sensitivity threshold triggered by 72 EtOH withdrawal was prevented with a MAGL inhibitor, and this effect was blocked by both CB_1_R and CB_2_R antagonists. To further characterize the specific role of 2-AG signaling in the regulation of mechanical hypersensitivity during EtOH withdrawal, the DAGL inhibitor DO34 was used to block the endogenous synthesis of 2-AG (Fig. 6A). After 3 weeks 2BC drinking, EtOH bottles were removed and the Von Frey testing was completed after 24 hours, 72 hours, and one week of withdrawal. DO34 pretreatment led to a robust emergence of previously unseen mechanical hypersensitivity at 24 hours withdrawal, with significantly lowered Von Frey thresholds (Fig. 6B). Next, as predicted, the mechanical thresholds of both vehicle and DO34 groups was very low on the Von Frey test at 72 hours withdrawal. Finally, DO34 completely prevented recovery of normal mechanical responses, and caused persistent withdrawal-triggered hypersensitivity at one-week into EtOH withdrawal (Fig. 6B-C). Alcohol consumption and preference for this cohort of mice is shown in Fig. 6D-E. Lastly, we showed that DO34 did not affect mechanical sensitivity thresholds in water drinking mice (Fig. 6F). These data indicate that inhibition of DAGL leads to an earlier onset and persistent mechanical hypersensitivity induced by EtOH withdrawal, and that the effects of 2-AG depletion are specific to the alcohol withdrawal state as DO34 has no effect on sensitivity thresholds in control mice.

**Figure 6.**
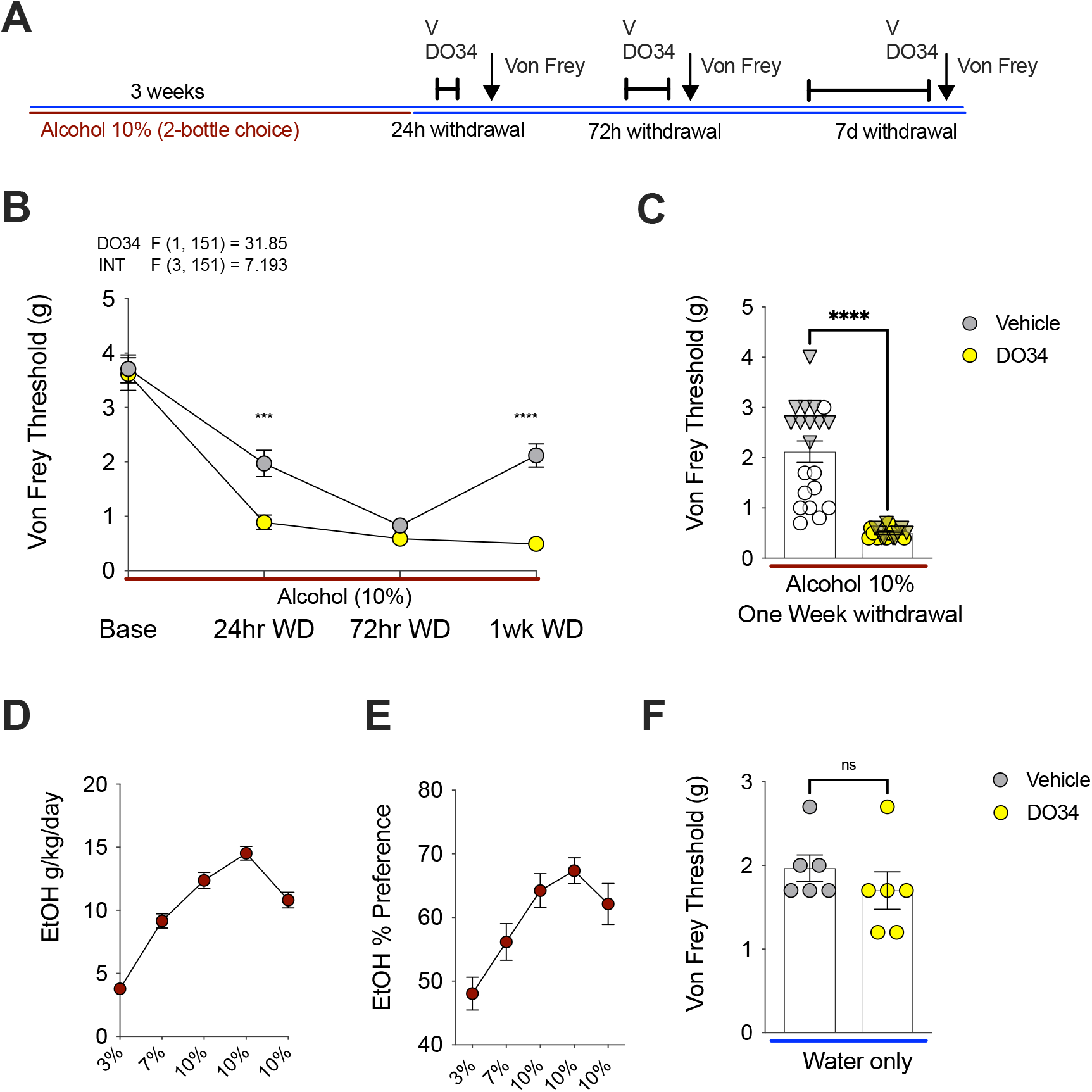
DO34 exacerbates mechanical hypersensitivity during EtOH withdrawal. **(A)** Schematic of experimental timeline and drug treatments. **(B)** DO34 decreased Von Frey threshold at 24-hours and one-week into EtOH withdrawal relative to vehicle-treated mice. **(C)** The same results at one week withdrawal separating males (circles) and female (triangles) data points. **(D-E)** Weekly EtOH intake and preference for mice shown in (B-C). (F) D034 does not affect sensitivity thresholds in water-only drinking mice. F-scores on graphs represent main effects and interaction from Two-way ANOVA. ***p<0.001, ****p<0.0001 via post-hoc Holm-Sidak pairwise comparisons or unpaired t-test. NS; not significant. Data presented as individual mice (triangles males and circles females) and Mean ± SEM.

## DISCUSSION

Here we show that pharmacological inhibition of MAGL prevents alcohol withdrawal-induced mechanical hypersensitivity via activation of CB_1_ and CB_2_ receptors. Importantly, neither 2-AG augmentation nor 2-AG inhibition affected mechanical sensitivity thresholds in water drinking mice or in mice exposed to alcohol but not in withdrawal. These data are partially consistent with recent data indicating MAGL inhibition within the lateral habenula reduces mechanical hypersensitivity rats undergoing alcohol withdrawal; however, in this study MAGL inhibition also increased mechanical thresholds to a similar degree in control rats [20]. We also showed that endogenous 2-AG serves to counteract alcohol withdrawal associated mechanical hypersensitivity, as blockade of 2-AG synthesis accelerated the onset and dramatically extended the time course of hyperalgesia in our model. That blockade of 2-AG synthesis had no effect on mechanical sensitivity thresholds in control mice suggests alcohol withdrawal is associated with an adapted state during which endogenous 2-AG exerts analgesic effects to limit the adverse consequences of withdrawal on pain sensitivity. These data demonstrate a critical role for 2-AG in the bidirectional modulation of mechanical sensitivity thresholds selectively during alcohol withdrawal and suggest MAGL inhibition could represent a novel approach to the treatment of hyperalgesic states associated with alcohol withdrawal.

MAGL inhibition has been shown to reduce neuropathic pain [75], pain induced by trigeminal nerve injury [76], chemotherapy-induced neuropathic pain [77], pain induced by models of multiple sclerosis [59], pain arising from inflammatory states [60], joint pain in osteoarthritis models [78], and chronic stress-induced hyperalgesia [79]. Previous studies have also implicated CB_1_ and CB_2_ receptors in the analgesic effects of MAGL inhibition in various preclinical models [58, 66, 75, 77]. Our data and a recent study [20] confirm the preclinical efficacy of MAGL inhibition in the reduction of mechanical hypersensitivity associated with alcohol withdrawal. There is some evidence that MAGL inhibition can increase hedonic behavior [80, 81], however we have previously shown that MAGL inhibition did not increase alcohol consumption or preference in mice [82], and direct lateral habenula MAGL inhibition actually reduced operant alcohol self-administration [20]. Moreover, MAGL inhibition has been shown to reduce negative affective states induced by alcohol withdrawal [83]. These findings, combined with our present results suggest MAGL inhibition could represent an important new approach to reduce negative reinforcement driven alcohol use by reducing both negative affective and somatic states associated with alcohol withdrawal without propensity to increase alcohol intake *per se*.

Augmentation of 2-AG levels via pharmacological MAGL inhibition has been shown to act on both CB_1_ and CB_2_ receptors to reduce pain sensitivity [58, 66, 75, 77, 84]. In the brain, the CB_1_ receptor is the most dominant eCB binding site [85, 86], while the CB_2_ receptor is largely associated with immune cells and microglia in the CNS [86–89]. Our pharmacological studies demonstrate that blockade of either CB_1_ receptors with Rimonabant or CB_2_ receptors with AM630 both prevent the anti-hyperalgesic effects of JZL184. These data indicate JZL184 given during alcohol withdrawal reduces mechanical hypersensitivity through activation of both CB_1_ and CB_2_ receptors, consistent with a dual receptor mechanism in a variety of other models noted above. The relative contribution of central vs. peripheral cannabinoid receptors to the anti-hyperalgesic effects of JZL184 during alcohol withdrawal is not known but is an important area for future investigation that could have important implications for 2-AG-based therapeutics development.

The present studies also support the critical role of endogenous 2-AG signaling in pain processing during alcohol withdrawal. Postsynaptic 2-AG is synthesized on-demand by the enzyme DAGLα in neurons [90–92]. Pharmacological inhibition of DAGL via the compound DO34 results in dramatic decreases in brain 2-AG levels [82, 93]. DAGLα^-/-^ mice exhibit more anxiety-like and depression-like behaviors at baseline [94, 95], and DO34 treatment after acute stress causes increased anxiety and depression-like behaviors [96]. Our lab has also shown DO34 impairs fear extinction in mice [97], and reduces alcohol consumption in a two-bottle choice paradigm [82]. However, little is known about the effects of inhibition of DAGL on pain processing. The current studies demonstrate that blocking synthesis of 2-AG exacerbates pain sensitivity, causing an earlier onset of mechanical hypersensitivity during withdrawal (24 hours), which persisted beyond the recovery time established in the previous experiments. Under the current experimental conditions, mice treated with DO34 demonstrated robust mechanical hypersensitivity that persisted 7 days into alcohol withdrawal. The current results are difficult to directly compare to previous findings, with one published study by Wilkerson and colleagues indicating pain sensitivity was reduced by DO34 in a model of LPS-induced allodynia [98]. However, this could be due in part to an anti-inflammatory effects secondary to reduced production of 2-AG-derived pro-inflammatory prostaglandins. We have shown previously that 2-AG derived proinflammatory prostaglandin glycerol esters (PG-Gs) can be generated following LPS-driven upregulation of cyclooxygenase-2 [99], and several other reports have shown LPS increases the production of pro-inflammatory prostaglandins [100–102]. In support of our current findings of reduced mechanical sensitivity thresholds in alcohol withdrawal after DO34 treatment, another study demonstrated that pharmacological inhibition of DAGL reduced orbital, but not hind paw, mechanical sensitivity thresholds [103]. Future studies will be needed to determine the specific conditions under which 2-AG signaling represents an anti-hyperalgesic system.

Previous studies have implicated CB_1_ receptors in the suppression of nociceptive transmission [104–108]. For example, Rimonabant administration results in hyperalgesia in the formalin paw test [109, 110] and hot plate test [111], and reverses stress induced analgesia [112]. Taken together, these data suggest alcohol withdrawal recruits 2-AG signaling to counteract mechanical hypersensitivity over extended time scales (at least ~1 week). Future studies aimed at elucidating the receptor mechanisms subserving this adaptive process would be of high significance and may reveal mechanistic insight into how 2-AG signaling regulates nociceptive processing as a function of alcohol withdrawal states.

We have previously shown that alcohol withdrawal increases negative affective states during protracted withdrawal in mice, some of which can be reversed by MAGL inhibition [113]. However, we did not detect clear changes in anxiety-like behaviors at the 72h time point in any assay we conducted, suggesting that the increase in mechanical sensitivity was not secondary to increases in anxiety-like states of mice. In contrast to our findings, mice treated with alcohol five consecutive days a week for 3-4 weeks via oral gavage exhibited both anxiety-like behavior and mechanical allodynia at 24 hours of alcohol withdrawal [15, 114]. However, this could be partially due to increased stress response seen with oral gavage methods [115, 116]. Others have reported the emergence of anxiety-like behaviors at different withdrawal time points, such as 5-50 hours into withdrawal, many using alcohol vapor exposure [117–127]. The findings are not easily compared between studies using different routes of administration and time of testing in withdrawal. Overall, the data presented here agree with literature showing three days as the optimal withdrawal time for induction hyperalgesia in mice following chronic alcohol drinking in a two-bottle choice model [128, 129]. Under the current experimental conditions of 72-hour withdrawal from alcohol, the effects on mechanical sensitivity and anxiety-like behaviors appear dissociable.

In clinical literature, irritability is one of three conditions contributing to a negative emotional states during drug withdrawal (dysphoria, anxiety, and irritability) [130, 131] and contributing to relapse [132]. Irritability-like behavior is challenging to model in rodents, with early alcohol withdrawal experiments relying on subjective experimenter observations of “irritability” in rodents [133–136]. The bottle brush test used here is a measure of irritability in rodents based on a tactile stimulus, which has been shown to detected the presence of irritabilitylike behavior during alcohol withdrawal [137] and oxycodone withdrawal [138]. It is possible that the decreased withdrawal thresholds (increased mechanical sensitivity) measured by the Von Frey tests could be related to irritability rather than hyperalgesia *per se*. Indeed, we did detect increases in irritability-like behavior at 72h of withdrawal, however, this effect was not reversed by JZL184 treatment. These data suggest that reduced withdrawal thresholds observed using the Von Frey test were not likely secondary to changes in irritability, since they were robustly reversed by JZL184, whereas irritability-like behaviors were not affected by JZL184.

While the current set of experiments focus on the eCB 2-AG, future work should determine how the eCB anandamide (AEA) may influence pain or hypersensitivity during alcohol withdrawal. Inhibition of fatty acid amide hydrolase (FAAH), the primary AEA degrading enzyme, abolished alcohol withdrawal anxiety seen following a single alcohol IP injection [139], and FAAH ^-/-^ mice are less susceptible to handling-induced convulsions following chronic ethanol exposure [140]. Furthermore, FAAH inhibition reportedly reduced allodynia when co-administered with a COX-2 inhibitor in models of neuropathy and inflammation [141], and FAAH inhibitor URB597 was analgesic in both control and ethanol withdrawal mice when injected into the lateral habenula [20]. Finally, in addition to MAGL, the ABHD6 enzyme hydrolyzes 2-AG in the brain and is located on postsynaptic neuronal elements [142]. The literature on the role of this enzyme in pain processing is limited, but there is some evidence inhibition of ABHD6 is antinociceptive in neuropathic pain models [143], and may regulate production of 2-AG derived pro-inflammatory prostaglandins [144]. Future studies examining these approaches would be important.

In summary, the present experiments provide insight into the regulatory role of 2-AG signaling in mechanical hypersensitivity during alcohol withdrawal. Inhibition of MAGL reduced mechanical hypersensitivity during alcohol withdrawal, while blocking 2-AG synthesis caused an earlier induction of hypersensitivity during withdrawal, and prevented recovery, with effects persisting up to 7 days into withdrawal. These data provide insight into the therapeutic potential of MAGL inhibitors in AUD.

## ACKNOWLEDGEMENTS

These studies were supported by NIH grants AA026186 (S.P.). The content is solely the responsibility of the authors and does not necessarily represent the official views of the National Institutes of Health.

